# Protein-protein docking with large-scale backbone flexibility using coarse-grained Monte-Carlo simulations

**DOI:** 10.1101/2021.02.22.432196

**Authors:** Mateusz Kurcinski, Sebastian Kmiecik, Mateusz Zalewski, Andrzej Kolinski

## Abstract

Most of the protein-protein docking methods treat proteins as almost rigid objects. Only the side-chains flexibility is usually taken into account. The few approaches enabling docking with a flexible backbone typically work in two steps, in which the search for protein-protein orientations and structure flexibility are simulated separately. In this work, we propose a new straightforward approach for docking sampling. It consists of a single simulation step during which a protein undergoes large-scale backbone rearrangements, rotations, and translations. Simultaneously, the other protein exhibits small backbone fluctuations. Such extensive sampling was possible using the CABS coarse-grained protein model and Replica Exchange Monte Carlo dynamics at a reasonable computational cost. In our proof-of-concept simulations of 62 protein-protein complexes, we obtained acceptable quality models for a significant number of cases.

## Introduction

Protein-protein interactions are fundamental in many biological processes. Their structural characterization is one of the biggest challenges of computational biology. A variety of docking methods are currently available for structure prediction of protein-protein complexes [1,2]. They can be divided into free (global) and template-based docking. Free (global) docking methods are designed to generate many distinct binding configurations. Template-based methods restrict docking to a binding mode found in a structural template. As demonstrated in the blind docking challenge, Critical Assessment of PRediction of Interactions (CAPRI), template-based methods generate more accurate results but only if a good quality template exists [1–5]. In some cases lacking useful templates, free global docking can yield acceptable results. According to recent estimates, the best free docking methods find adequate models among the top 10 predictions for around 40% of the targets [1]. The CAPRI analysis also indicates that protein backbone flexibility is a big challenge; protein complexes that undergo substantial conformational changes upon docking get no successful predictions from any method [3–5].

Presently, most of the free docking methods treat the backbone of input protein structures as rigid. This approximation reduces the protein-protein docking problem to a 6D (three rotational and three translational degrees of freedom) search space. Rigid-body search for the binding site most often rely on the Fast Fourier Transform [6–8]. Other successful approaches include 3D Zernike descriptor-based docking [9,10] or geometric hashing [11]. These rigid-body methods are often used as a first docking step, followed by scoring [12–16], using experimental data [17] and/or structural refinement to capture backbone flexibility [5,18]. Molecular Dynamics is perhaps the most common refinement strategy, either in classic or enhanced sampling versions [17,19–22]. Other tools use rotamer libraries to address side-chain flexibility [23] and Elastic Network Models (ENM) for modeling backbone rearrangements [24–28]. Accounting for backbone flexibility in the search for the binding site significantly increases the docking complexity and makes it practically intractable using conventional all-atom modeling approaches. This enormous computational complexity of flexible docking can be reduced using coarse-grained protein models [29–32]. The best-performing methods that can now include backbone flexibility during the docking calculations use coarse-grained models and/or ENM-driven simulations. These include RosettaDock combining coarse-grained generation of backbone ensembles and all-atom refinement [33–35]; ATTRACT combining coarse-grained docking with ENM and all-atom refinement [36,37]; and SwarmDock using all-atom ENM [25,38]. All these approaches show some advantages in modeling protein flexibility compared to rigid-body docking followed by structure refinements. However, effective modeling flexibility in protein-protein docking remains an unsolved problem, as demonstrated in the recent CAPRI round [25,35,37,39].

In this work, we use a well-established CABS coarse-grained protein model [29] for protein-protein docking. During the CABS docking simulation, one of the docking partners undergoes a long random process of rotations, translations, and extensive backbone conformational rearrangements that significantly modify its fold. Simultaneously, the backbone of the second protein undergoes small fluctuations.

## Methods

### Docking simulation protocol

In this work, we present the protein-protein docking simulation protocol that relies on the CABS coarse-grained model. The CABS design and applications have been recently described in the reviews on protein coarse-grained [29] and protein flexibility [24,40] modeling. Here we outline only its main features. The CABS model uses a coarse-grained representation of protein chains (see Figure 1), Replica Exchange Monte Carlo (REMC) dynamics, and knowledge-based statistical potentials. Representation of protein chains is based on C-alpha traces, restricted to an underlying high-resolution lattice. The lattice spacing allows slight fluctuations of the C-alpha – C-alpha distances and many pseudo-bonds orientations. Virtual pseudo-atoms are placed in the centers of these C-alpha–C-alpha bonds and are used to locate the main-chain hydrogen bonds. Also, the positions of the two pseudo-atoms representing side chains are defined by the geometry of C-alpha traces and amino-acid identities. Such fixed positions of side chains (taken from the statistics of protein databases) reduce the model’s resolution. However, this limitation is less serious than it may appear since even small movements of the main chain (allowed due to the soft nature of the assumed geometrical restrictions) leads to large moves of the side chains. This way, the packing of side chains can be quite accurate. The interaction scheme of CABS consists of statistical potentials mimicking effects of main chain rotational preferences, main-chain hydrogen bonds, and side-chain contacts. All statistical potentials, derived from structural regularities observed in PDB structures, have relatively broad minima compensating the low-resolution effects and allowing a fast search for global energy minima. The solvent is treated implicitly, and its averaged effects are encoded within the above-mentioned contact potentials. Energy computation for protein chain models is very fast since many interactions could be pre-computed (and coded in large tables) due to the discretized patterns of main chains geometry. The Monte Carlo sampling of CABS uses a set of local movers. The resulting model dynamics is quite realistic for large-scale distances, allowing coarse-grained modeling of protein structures, dynamics, and protein-protein interactions.

**Figure 1.**
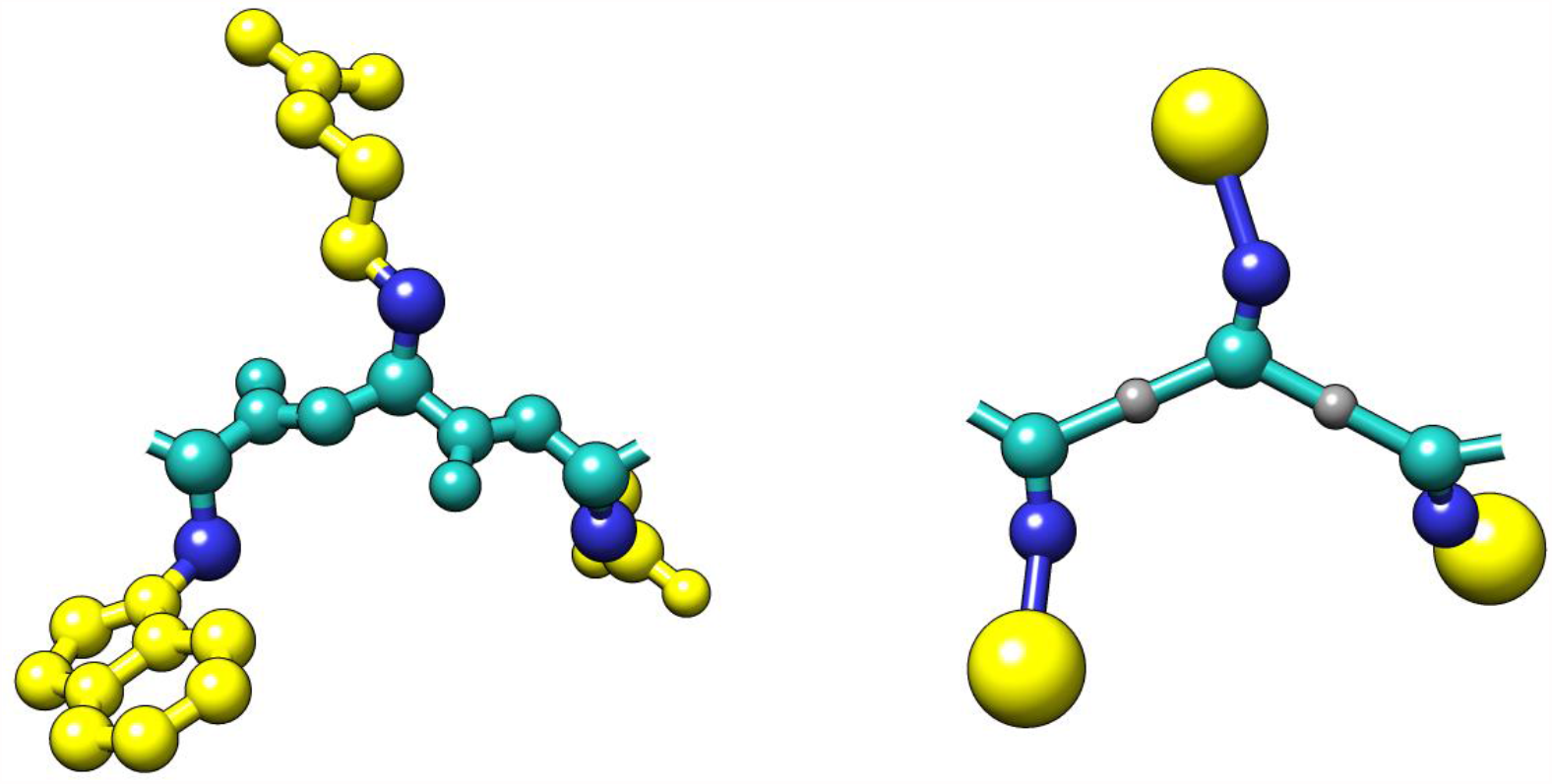
Comparison of the all-atom (left) and the CABS coarse-grained model representation (right) for an example tripeptide. In the CABS model, protein residues are represented using C-alpha, C-beta, united side-chain atom, and the peptide bond center [29].

The modeling protocol consists of the following steps:

i. **Preparing input structures of a protein-ligand and a protein-receptor**. The protocol requires the input of two protein structures (single-or multi-chain) in the PDB format. One of them has to be indicated as a ligand and the second as a receptor. The ligand undergoes large conformational fluctuations, translations, and rotations around the receptor within the proposed protocol. The ligand should be a smaller protein because the computational cost of searching its conformational space rapidly grows with the chain length. That is because the motion of the entire structure (including fold relaxation, rotation, and translation of the entire molecule) is simulated by a random sequence of local moves. The accuracy of such sampling is acceptable for not too-large proteins. On the other hand, treating the ligand-protein as a fully flexible object allows approximate studies of entire docking trajectories. In some cases, it would be perhaps worth treating a larger protein (but not too large) as a flexible ligand, although this was out of range of the present studies.
ii. **Generating starting structures** Starting conformations are built using C-alpha coordinates only (in the CABS model C-alpha traces define position of other united pseudo-atoms, see details [29]). The algorithm places the protein-ligand center at 20 random positions around the protein receptor at the approximate distance of 20 Å from the protein receptor’s surface. Next, these protein-ligand systems are used as starting conformations for the 20 replicas in the REMC CABS sampling scheme (each replica starts from a different ligand-receptor arrangement).
iii. **Docking simulations using CABS coarse-grained model and REMC dynamics.** During simulations, the protein receptor structure is kept close to the starting structure using distance restraints. Distance restraints are generated using the input coordinates of the C-alpha atoms. Two residues are automatically restrained if two conditions are met. First, their separation along the sequence has to be at least five residues. Second, the distance between their C-alpha atoms must be within the range of 5–15 Å. During simulations, the receptor restraints imply small-scale fluctuations of the protein receptor backbone in the range of 1 Å and, accordingly, more significant fluctuations of the side-chain atoms. A similar restraints scheme is applied to the protein-ligand but with tenfold weaker weights. During simulations, the ligand moves freely within the vicinity of the receptor and internal restraint allows for large-scale fluctuations of its structure. Usually, the ligand fluctuations are within the range of 2 and 12 Å to the input structure although folding-unfolding events are possible at highest temperatures. The docking simulation is conducted using CABS REMC pseudo-dynamics with simulated annealing. In this work, 20 replicas and 20 annealing steps have been used. All the REMC scheme parameters have been adjusted to allow for large-scale conformational transitions, rotations, and translations of the protein-ligand in a reasonable computational time. The modeling protocol collects trajectories from all 20 replicas. The protocol saves only a small fraction (2%) of the generated models for further analysis i.e., 500 models from each replica, thus 10,000 models in total.
iv. **Reconstruction of the CABS model representation.**The set of 10,000 models stored as C-alpha traces are reconstructed to complete CABS model representation using CABS algorithm [29]. In CABS, positions of C-beta and Side-Chain united atoms are defined by the positions of the three consecutive C-alpha atoms and the amino acid identity (the most probable positions from the PDB database are used).
v. **Clustering of contact maps.** First, for all of the 10’000 models the contact maps between the receptor and the ligand proteins are calculated. Two residues are considered to form a contact if their Side Chain pseudoatoms are at most 6 Å apart (for Alanines the C-beta atoms are considered as the Side Chain; for Glycines – it’s the C-alpha atoms). Next, the algorithm sorts the models according to the number of the receptor-ligand contacts, and the set of top 1000 is kept for further processing. This way the transient and weakly bound complexes are removed from the solutions pool. In the next step, the 1000 contact maps are clustered together to identify the most frequently occurring contact patterns. The complete link hierarchical clustering was used with the Jaccard index as the distance metric between contact maps. Finally, the identified clusters are ranked according to their density, defined as the number of the cluster members divided by the average metric between them.
vi. **Reconstruction to all-atom representation.** Representative models from the ten most dense clusters are reconstructed to all-atom representation using Modeller-based rebuilding procedure [41] (or can be reconstructed using other rebuilding strategies, see review [42]).

In recent years, the CABS model has been used for modeling the flexibility of globular proteins [43–46] and various processes leading to large-scale conformational transitions. These included: ab initio simulations of protein folding mechanisms [47,48], folding and binding mechanisms [49,50], and free protein-peptide docking within the CABS-dock tool [51–57]. The CABS-dock is a well-established peptide docking tool that has been made available as a web server [51,52] and, most recently, as a standalone application [54]. Its distinctive feature among other tools is the possibility of fast simulation of the large backbone rearrangements of both peptide and protein receptors during binding (see the review on protein-peptide docking tools [58]). In addition, the CABS-dock has been used in multiple applications (recently reviewed [56]), including docking to receptors with disordered fragments [40,50], GPCRs [59], and modeling proteolysis mechanisms [60].

The presented protocol for protein-protein docking utilizes the CABS-dock standalone package [54] developed primarily for protein-peptide docking. In order to tackle the protein-protein docking problem, key changes have been made to the docking algorithm that aimed mainly at the improvement of the conformational sampling. First of all, the temperature distribution between replicas in the REMC scheme was adjusted. Instead of constant temperature increment between consecutive replicas, as in the original CABS-dock, here we’ve implemented progressive geometric raise of the temperature increment. Furthermore, the number of simulation replicas was increased to twenty versus ten in the original CABS-dock. Besides the sampling improvement, a new clustering protocol was introduced. The original CABS-dock used RMSD-based clustering, which worked well for peptides. For the protein-protein complexes, however, purely geometrical similarity condition such as the RMSD is too severe. Namely, for two binding poses, where the mobile protein was docked in the exact same pocket but is slightly tilted in one of them, the RMSD difference would be considerable. Despite representing similar binding poses, the two structures would end up in different clusters. To overcome this, the current protocol uses clustering based on the similarity between receptor-ligand contact maps.

### Results analysis and quality metrics

The docking simulation analysis was performed using Python and NumPy (Python library). Structural differences between experimentally determined structures and generated models were evaluated using Root Mean Square Deviations (RMSDs). Interface RMSD (iRMSD) is an RMSD calculated for interface residues of the receptor and the ligand separated by no more than 6 Angstroms. Ligand RMSD (LRMSD) is an RMSD computed for the ligands after the superimposition of the receptors. Ligand only RMSD (LoRMSD) is an RMSD computed for the ligand structure only. Root Mean Square Fluctuation (RMSF) is a measure of the amino acid’s flexibility. It is calculated for every residue as the square root of this residue’s variance around the reference residue position. The fraction of native contacts (fNAT) was calculated as a number of experimental structure contacts found in the generated structure divided by the total number of contacts found in the experimental structure. Rather restrictive criterion of a contact, distance up to 6 Å between side chain centers, was used. All figures presented in this work were generated using PyMOL, UCSF Chimera, and Matplotlib (Python library).

### Dataset

In this docking study, we used protein-protein cases from the ZDOCK benchmark set[61] (cases in which a smaller size protein – a protein ligand – contained more than one protein chain, or chain gaps, were discarded from our set). The set comprises the three flexibility-based subsets: low-flexible (almost rigid), medium-flexible and highly-flexible with available unbound X-ray structures of both the protein-receptor and the protein-ligand. The unbound structures were used as the docking input. As the reference for calculating various similarity measures, we used the X-ray structures of the protein-ligand complexes. Table 1 lists all the PDB IDs of X-ray structures used in the study.

**Table 1.** Summary of the docking simulations. The table characterizes X-ray data used in the docking, average ligand flexibility during simulations (average LoRMSD from 10’000 models), and docking results. The table reports the best accuracy models out from all (10’000) and 10 top-scored models. The metrics definitions are provided in the Methods section. The table divides the presented cases on the three categories: low-flexibility, medium-flexibility and highly flexible cases.

| X-ray data<br>(number of residues) |  |  |  | Ligand<br>flexibility | Results - best<br>from all models |  |  | Results - best<br>from 10 top-scored<br>models |  |  |
| --- | --- | --- | --- | --- | --- | --- | --- | --- | --- | --- |
| Receptor* | Ligand* | Complex | RMSD** | Average<br>LoRMSD | iRMSD | LRMSD | fNAT | iRMSD | LRMSD | fNAT |
| Low-flexibility cases |  |  |  |  |  |  |  |  |  |  |
| 5CHA<br>(238) | 2OVO<br>(53) | 1CHO | 0.62 | **4.84** | **2.65** | 6.95 | 0.48 | 2.96 | 10.93 | 0.18 |
| 2PKA<br>(232) | 6PTI<br>(56) | 2KAI | 0.91 | **4.64** | **3.32** | 11.34 | 0.19 | 4.75 | 15.76 | 0.12 |
| 1CHG<br>(245) | 1HPT<br>(56) | 1CGI | 1.53 | **5.24** | **2.76** | 4.13 | 0.37 | 6.18 | 14.15 | 0.09 |
| 2PTN<br>(223) | 6PTI<br>(58) | 2PTC | 0.31 | **5.23** | **2.97** | 11.86 | 0.29 | 4.39 | 15.93 | 0.15 |
| 1SUP<br>(275) | 2CI2<br>(64) | 2SNI | 0.37 | **3.89** | **1.09** | 3.86 | 0.69 | 2.81 | 9.09 | 0.46 |
| 2ACE<br>(532) | 1FSC<br>(61) | 1FSS | 0.76 | **4.48** | **3.41** | 7.20 | 0.25 | 15.03 | 32.56 | 0.03 |
| 1MAA<br>(533) | 1FSC<br>(61) | 1MAH | 0.60 | **4.58** | **2.49** | 3.89 | 0.45 | 11.25 | 24.43 | 0.06 |
| 1A2P<br>(108) | 1A19<br>(89) | 1BRS | 0.47 | **3.33** | **1.94** | 4.19 | 0.64 | 4.01 | 8.74 | 0.14 |
| 1CCP<br>(294) | 1YCC<br>(103) | 2PCC | 0.39 | **4.18** | **3.13** | 10.19 | 0.25 | 11.89 | 26.68 | 0.08 |
| 1SUP<br>(275) | 3SSI<br>(107) | 2SIC | 0.39 | **4.01** | **4.03** | 18.96 | 0.23 | 4.77 | 19.40 | 0.12 |
| 1VFA<br>(223) | 1LZA<br>(129) | 1VFB | 0.59 | **3.72** | **4.61** | 15.07 | 0.11 | 17.45 | 37.15 | 0.00 |
| 1MLB<br>(432) | 1LZA<br>(129) | 1MLC | 0.85 | **3.74** | **2.82** | 10.47 | 0.36 | 8.04 | 33.09 | 0.04 |
| Medium-flexibility cases |  |  |  |  |  |  |  |  |  |  |
| 1CHG<br>(226) | 1HPT<br>(56) | 1CGI | 2.02 | 5.80 | **2.46** | 3.21 | 0.44 | 5.86 | 10.72 | 0.12 |
| 5C2B<br>(241) | 4ZAI<br>(80) | 5CBA | 1.49 | 4.51 | **2.48** | 7.64 | 0.42 | 9.34 | 16.01 | 0.10 |
| 5P21<br>(166) | 1LXD<br>(87) | 1LFD | 1.79 | 4.12 | **2.87** | 6.76 | 0.27 | 12.47 | 24.24 | 0.00 |
| 1R6C<br>(142) | 2W9R<br>(97) | 1R6Q | 1.67 | 9.27 | **7.95** | 11.97 | 0.14 | 13.71 | 35.71 | 0.00 |
| 1JXQ<br>(242) | 2OPY<br>(106) | 1NW9 | 1.97 | 4.09 | **7.05** | 8.69 | 0.23 | 9.33 | 17.55 | 0.00 |
| 1IAS<br>(330) | 1D6O<br>(107) | 1B6C | 1.96 | 4.65 | **4.72** | 10.74 | 0.14 | 12.24 | 23.99 | 0.00 |
| 5E56<br>(116) | 5E03<br>(113) | 5E5M | 1.56 | 4.16 | **3.83** | 9.09 | 0.23 | 10.96 | 20.00 | 0.00 |
| 2HRA<br>(180) | 2HQT<br>(115) | 2HRK | 2.03 | 7.27 | **3.55** | 10.17 | 0.26 | 10.81 | 32.54 | 0.00 |
| 4BLM<br>(256) | 4M3J<br>(116) | 4M3K | 1.77 | 4.41 | **4.96** | 7.41 | 0.10 | 13.75 | 27.11 | 0.03 |
| 1E78<br>(578) | 5VNV<br>(120) | 5VNW | 1.49 | 3.81 | **5.93** | 22.23 | 0.10 | 23.89 | 70.83 | 0.00 |
| 3BX8<br>(167) | 3OSK<br>(121) | 3BX7 | 1.63 | 4.63 | **4.94** | 17.46 | 0.28 | 6.22 | 20.32 | 0.12 |
| 6ETL<br>(124) | 4POY<br>(121) | 4POU | 1.83 | 4.01 | **2.91** | 10.16 | 0.50 | 6.55 | 19.65 | 0.25 |
| 4FUD<br>(246) | 5HDO<br>(126) | 5HGG | 0.84 | 4.22 | **3.59** | 12.52 | 0.19 | 13.00 | 29.3 | 0.00 |
| 3TGR<br>(346) | 3R0M<br>(127) | 3RJQ | 0.79 | 4.00 | **5.32** | 16.98 | 0.13 | 12.77 | 33.94 | 0.00 |
| 6EY5<br>(585) | 5FWO<br>(129) | 6EY6 | 1.90 | 3.86 | **3.83** | 6.03 | 0.14 | 12.89 | 27.61 | 0.00 |
| 1SZ7<br>(159) | 2BJN<br>(141) | 2CFH | 1.55 | 5.13 | **1.98** | 4.01 | 0.71 | 2.82 | 5.50 | 0.63 |
| 3V6F<br>(437) | 3KXS<br>(142) | 3V6Z | 1.83 | 7.11 | **6.12** | 16.68 | 0.15 | 6.66 | 20.06 | 0.06 |
| 3CPI<br>(437) | 1G16<br>(156) | 3CPH | 2.12 | 4.34 | **4.87** | 15.64 | 0.09 | 15.02 | 27.88 | 0.00 |
| 1QJB<br>(460) | 1KUY<br>(166) | 1IB1 | 2.09 | 4.22 | **6.56** | 14.83 | 0.13 | 16.10 | 46.26 | 0.00 |
| 1IAM<br>(185) | 1MQ9<br>(173) | 1MQ8 | 1.76 | 4.22 | **4.93** | 14.99 | 0.21 | 26.17 | 70.50 | 0.00 |
| 3HI5<br>(430) | 1MJN<br>(179) | 3HI6 | 1.65 | 3.77 | **5.79** | 23.30 | 0.21 | 19.38 | 49.77 | 0.00 |
| 2G75<br>(429) | 2GHV<br>(183) | 2DD8 | 2.19 | 5.37 | **5.73** | 13.78 | 0.09 | 17.20 | 34.33 | 0.00 |
| 1A12<br>(401) | 1QG4<br>(202) | 1I2M | 2.12 | 4.19 | **2.84** | 6.43 | 0.51 | 3.58 | 6.97 | 0.47 |
| 1N0V<br>(825) | 1XK9<br>(204) | 1ZM4 | 2.11 | 3.54 | **8.82** | 28.17 | 0.04 | 11.05 | 48.14 | 0.00 |
| 4EBQ<br>(429) | 4E9O<br>(230) | 4ETQ | 0.47 | 3.72 | **7.12** | 14.74 | 0.20 | 8.68 | 19.61 | 0.07 |
| 1S3X<br>(380) | 1XQR<br>(259) | 1XQS | 1.77 | 5.44 | **5.63** | 26.14 | 0.11 | 15.88 | 30.51 | 0.00 |
| 3HEC<br>(329) | 3FYK<br>(282) | 2OZA | 1.89 | 4.29 | **4.35** | 9.32 | 0.33 | 11.24 | 18.8 | 0.03 |
| 6A0X<br>(437) | 2FK0<br>(322) | 6A0Z | 1.28 | 5.75 | **5.75** | 25.59 | 0.16 | 11.43 | 31.39 | 0.00 |
| 1RRP<br>(338) | 1YRG<br>(343) | 1K5D | 1.19 | 4.58 | **7.51** | 19.93 | 0.07 | 10.11 | 31.83 | 0.00 |
| **Highly flexible cases** |  |  |  |  |  |  |  |  |  |  |
| 1CL0<br>(316) | 2TIR<br>(108) | 1F6M | 4.9 | 3.83 | **7.02** | 11.34 | 0.10 | 11.92 | 18.06 | 0.00 |
| 1X9Y<br>(346) | 1NYC<br>(110) | 1PXV | 2.63 | 4.86 | **5.74** | 14.10 | 0.07 | 7.46 | 16.31 | 0.02 |
| 1JZO<br>(431) | 1JPE<br>(116) | 1JZD | 2.71 | 4.65 | **4.98** | 8.13 | 0.28 | 13.38 | 34.05 | 0.00 |
| 5D7S<br>(423) | 2GMF<br>(121) | 5C7X | 2.26 | 4.17 | **4.12** | 13.61 | 0.34 | 4.69 | 16.74 | 0.20 |
| 1FCH<br>(302) | 1C44<br>(123) | 2C0L | 2.62 | 5.51 | **5.02** | 5.54 | 0.21 | 10.24 | 24.74 | 0.00 |
| 1YWH<br>(268) | 2I9A<br>(123) | 2I9B | 3.79 | 7.14 | **5.79** | 17.59 | 0.14 | 6.92 | 33.23 | 0.05 |
| 3L88<br>(550) | 1CKL<br>(126) | 3L89 | 2.51 | 9.86 | **4.83** | 10.90 | 0.17 | 17.84 | 31.87 | 0.00 |
| 1ZM8<br>(239) | 1J57<br>(143) | 2O3B | 3.13 | 6.20 | **4.76** | 16.43 | 0.18 | 15.34 | 31.95 | 0.00 |
| 1G0Y<br>(310) | 1ILR<br>(145) | 1IRA | 8.38 | 4.07 | **12.97** | 22.24 | 0.08 | 15.86 | 25.46 | 0.05 |
| 1QUP<br>(219) | 2JCW<br>(153) | 1JK9 | 2.51 | 9.40 | **8.07** | 13.85 | 0.10 | 17.41 | 30.74 | 0.00 |
| 1SYQ<br>(259) | 3MYI<br>(163) | 1RKE | 4.25 | 4.15 | **5.26** | 6.43 | 0.38 | 16.11 | 34.67 | 0.00 |
| 2II0<br>(463) | 1CTQ<br>(166) | 1BKD | 2.86 | 4.51 | **4.80** | 7.33 | 0.14 | 19.96 | 39.32 | 0.00 |
| 1ERN<br>(416) | 1BUY<br>(166) | 1EER | 2.44 | 5.22 | **12.97** | 13.18 | 0.02 | 17.12 | 30.73 | 0.00 |
| 3AVE<br>(419) | 1FNL<br>(173) | 1E4K | 2.60 | 5.32 | **3.44** | 10.07 | 0.43 | 7.59 | 24.33 | 0.13 |
| 1R8M<br>(195) | 1HUR<br>(180) | 1R8S | 3.73 | 5.50 | **6.67** | 13.41 | 0.09 | 15.15 | 25.10 | 0.00 |
| 1QFK<br>(348) | 1TFH<br>(182) | 1FAK | 6.18 | 5.64 | **8.97** | 15.57 | 0.16 | 15.59 | 34.46 | 0.00 |
| 1F59<br>(440) | 1QG4<br>(202) | 1IBR | 2.54 | 5.01 | **6.65** | 14.36 | 0.14 | 16.41 | 33.07 | 0.00 |
| 4DVB<br>(427) | 4DVA<br>(246) | 4DW2 | 2.27 | 3.85 | **6.61** | 21.91 | 0.14 | 9.94 | 29.27 | 0.00 |
| 1NG1<br>(294) | 2IYL<br>(271) | 2J7P | 2.67 | 4.51 | **8.87** | 18.77 | 0.11 | 18.46 | 48.05 | 0.00 |
| 1UX5<br>(411) | 2FXU<br>(360) | 1Y64 | 4.69 | 4.15 | **6.42** | 13.50 | 0.27 | 15.50 | 36.42 | 0.00 |
| 1D0N<br>(729) | 1IJJ<br>(371) | 1H1V | 6.62 | 3.44 | **7.92** | 31.14 | 0.36 | 29.12 | 65.07 | 0.03 |

\* 4-letter PDB code for the crystal structures used in this study. \*\*The RMSD (in Å) of the interface Cα atoms for input receptor and ligand after superposition onto the co-crystallized complex system.

## Results

The most accurate models (out of the sets of 10’000 generated models and 10 top-scored) are characterized in Table 1. The Table presents different metrics of similarity to the experimental structures for the set of 62 protein-peptide complexes (divided into three categories: low, medium and high flexibility cases). To assess the sampling performance, below we will use the iRMSD values for the best models out of all models. According to the iRMSD values the CABS-based docking algorithm produced a significant number of near-native protein-protein arrangements of acceptable quality (iRMSD < 4 Å, according to CAPRI criteria) for most protein-protein cases in the categories of low and medium flexibility cases. However, in the high-flexibility category, the best iRMSD values are noticeably higher (in the range of 4-12 Angstroms). This results from the adopted distance restraint scheme (see the Methods section), which is uniform for whole proteins and introduces a penalty for deviations of more than 1 A from the input structures (unbound experimental structures). This penalty is very small for the protein ligands. Thus, the distance-restraints scheme allows for the large-scale conformational changes, however, it may also prevent from binding-induced conformational changes in the high-flexibility category. Therefore, there is the need to modify such a scheme for the most challenging targets.

The results analysis below focuses on the sampling performance for the selected low-flexibility barnase/barstar case. Figure 2 characterizes iRMSD versus CABS model energy values for the barnase/barstar (1BRS) and another low-flexibility case with clearly the lowest iRMSD value 1.09 Angstroms (2SNI).

**Figure 2.**
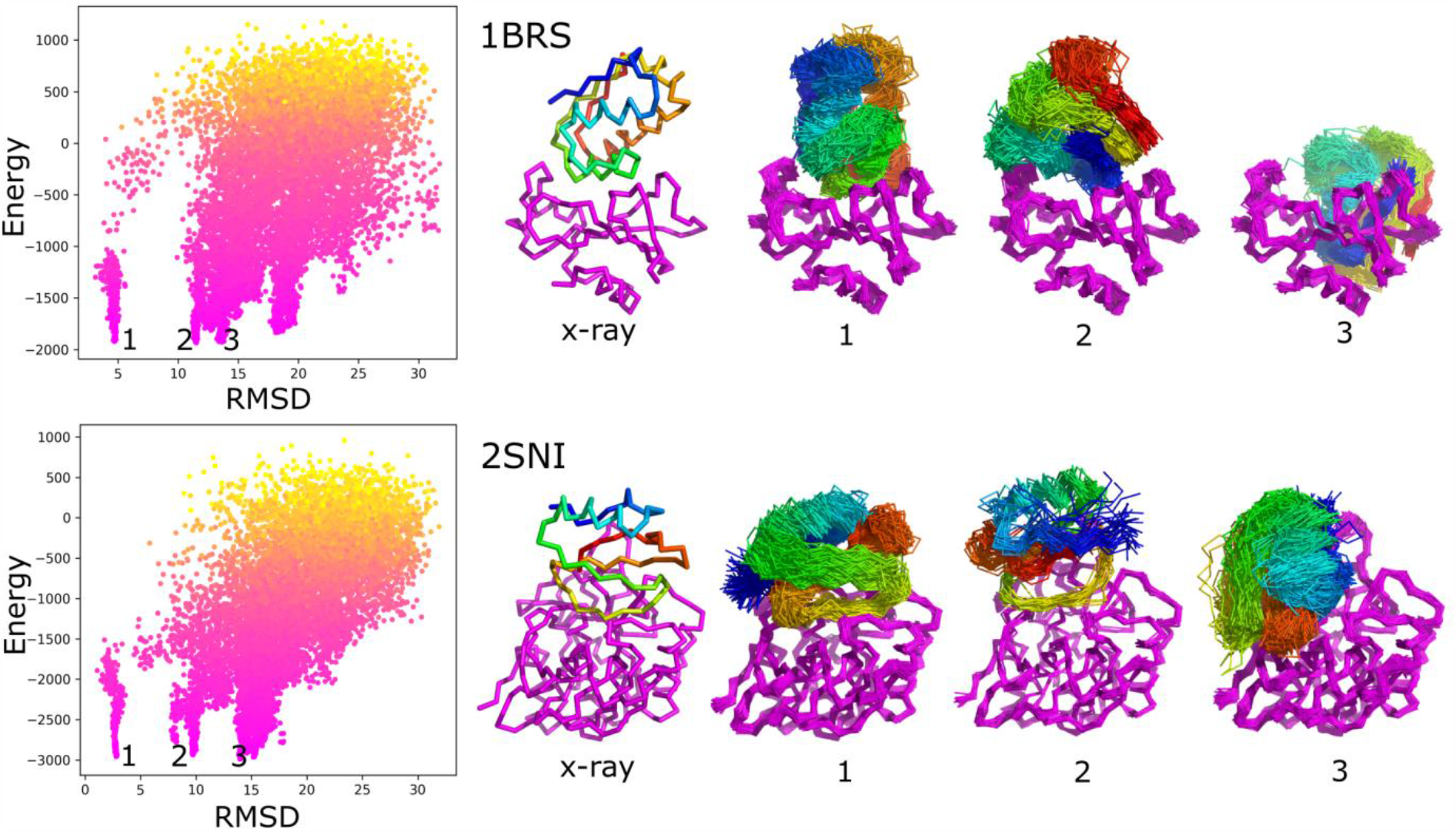
Characterization of docking results using RMSD to the X-ray structure and system energy. The left panels show the interface-RMSD versus CABS energy values. Point color represents the temperature – from yellow(high) to pink(low). The molecular visualizations show x-ray structures and ensembles of predicted models corresponding to selected energy minima (numbered in the picture from 1 to 3). As presented in the picture, the minima numbered as 1st corresponds to near-native protein-protein arrangements, others to non-native ensembles, as presented in the picture. The presented ensembles are the sets of similar models found in the structural clustering of contact maps (see Methods). The figure shows two modeling cases: 1BRS and 2SNI.

Figure 3 shows example ensembles of barnase/barstar models and the most accurate model (iRMSD 1.9 Å). A single system replica can explore an ample conformational space that involves significantly different binding configurations and protein-ligand conformations, as demonstrated in Figure 3c and Movie S1. Figure 4a further characterizes this single replica’s using iRMSD and LoRMSD (RMSD for ligand only) values. As presented in the figure, the ligand structure fluctuates around 5 Å (the same fluctuations in the context of all replicas are shown in Figure 4b). The ligand becomes significantly more closer to the X-ray structure after binding to the native binding site as reflected by iRMSD values. Namely, after correct binding, LoRMSD values get noticeably lower to around 2 Å (see Figure 4a). In the following sections, we do not discuss this aspect of our method; however, it is worth mentioning that the proposed method enables a detailed analysis of plausible docking trajectories. The described docking procedure uses REMC protocol enhanced by simulated annealing of all 20 replicas. Figure 4c shows their evolution through different temperatures. Figures 5 provide more detailed pictures of structural flexibility for protein “receptor” and “ligand”. Protein-protein contacts defining the complex assembly are characterized in Figure 6. In the presented example, the most persistent protein-protein contacts occur in about 15% of snapshots. Therefore, they are significantly less stable compared to intramolecular protein contacts (Figure 5).

**Figure 3.**
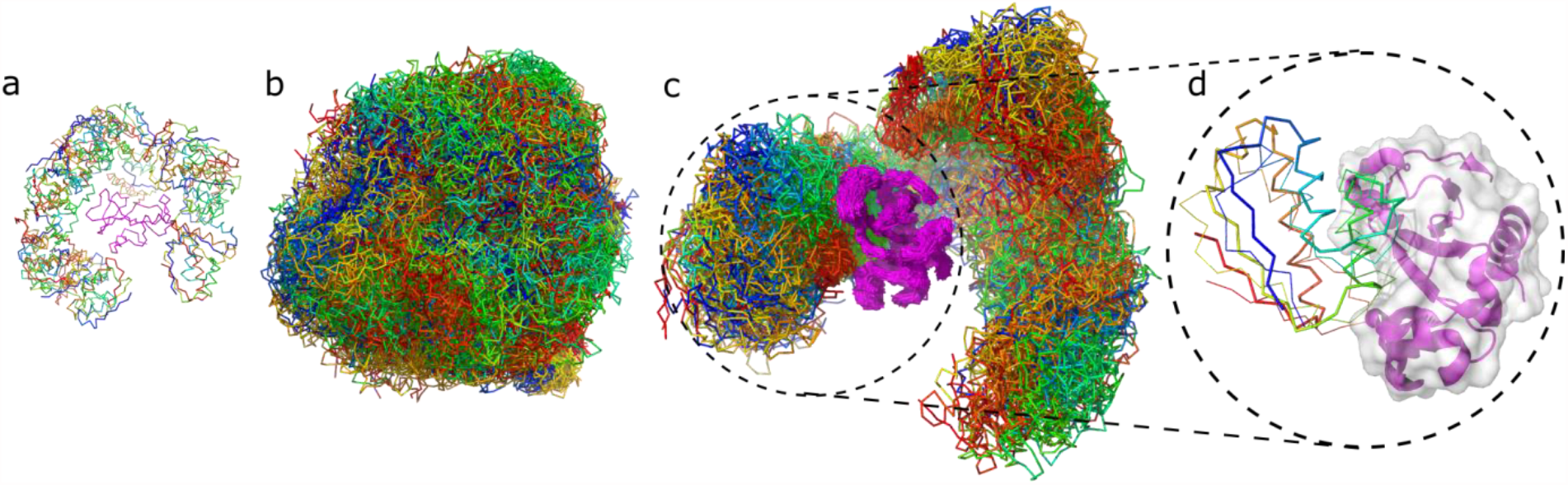
Protein-protein docking stages illustrated by barstar/barnase docking case. The figure shows the barnase receptor in magenta and the barstar ligand in rainbow colors. The respective panels show: (a) 20 starting structures for each replica of the system; (b) 10’000 models combined from 20 replicas (500 models per replica) in which the highly flexible ligand is covering the entire surface of the flexible receptor; (c) 500 models from 1 replica only, (d) the best model obtained for barnase/barstar system (the x-ray structure of the ligand is shown in thick ribbon, the modeled in thin ribbon).

**Figure 4.**
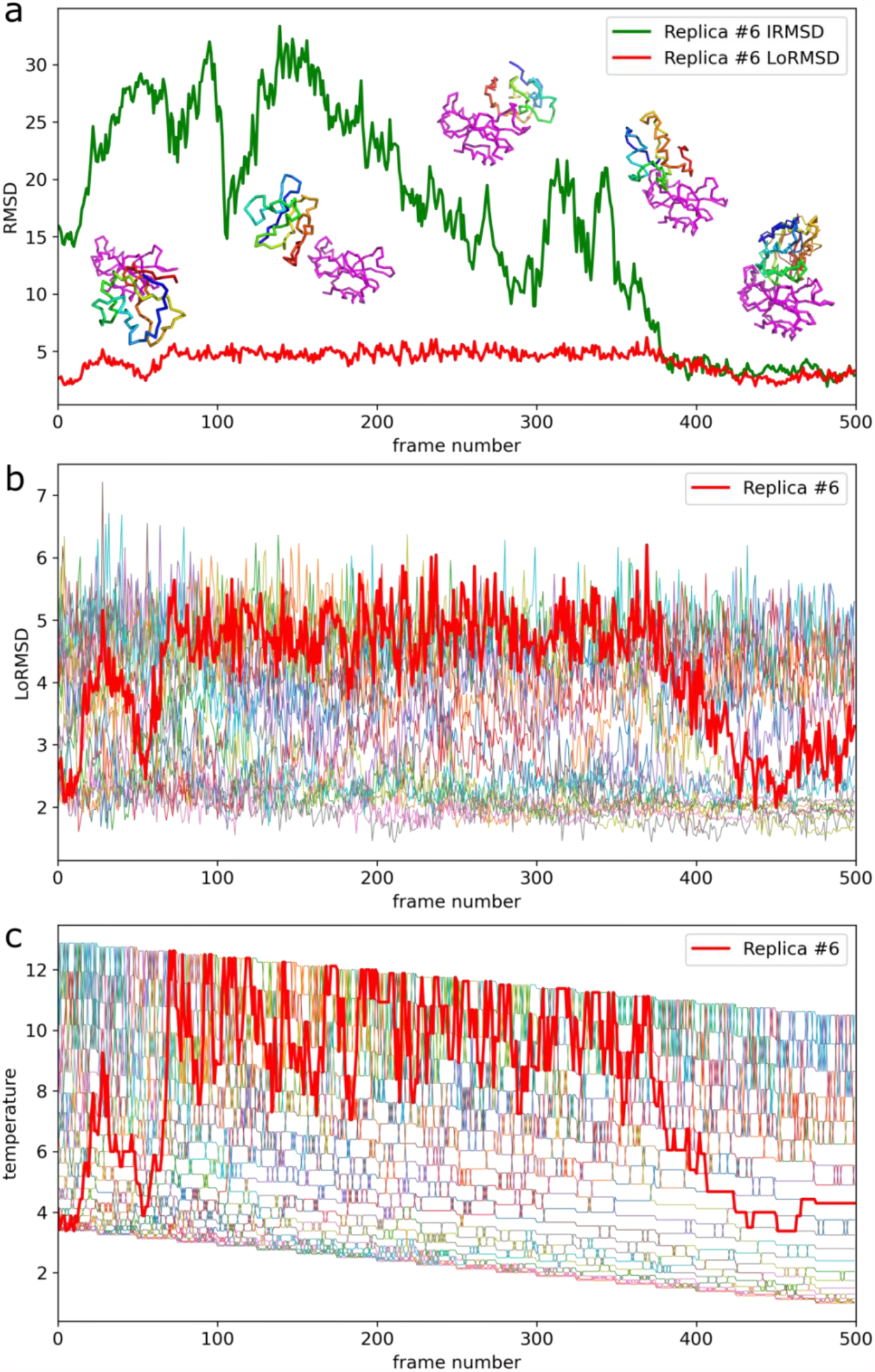
Docking trajectory for the selected replica of barnase/barstar system. The presented replica reached the most accurate barnase/barstar complex structure. (a) iRMSD (interface RMSD) and LoRMSD (ligand only RMSD) values. Example simulation snapshots illustrate the plot. The ligand is presented in rainbow colors, the receptor in magenta. The lowest iRMSD model (1.9 A from x-ray structure) is presented on the right lower corner superimposed on the x-ray structure (the x-ray structure is shown in thick lines, the predicted model in thin lines). (b) ligand only RMSD (LoRMSD) values for all replicas. The thick red line presents selected replica. (c) exchange of system replicas between different temperatures driven by Replica Exchange Monte Carlo (REMC) system. The thick red line presents selected replica. The replica trajectory is also presented in the Movie S1.

**Figure 5.**
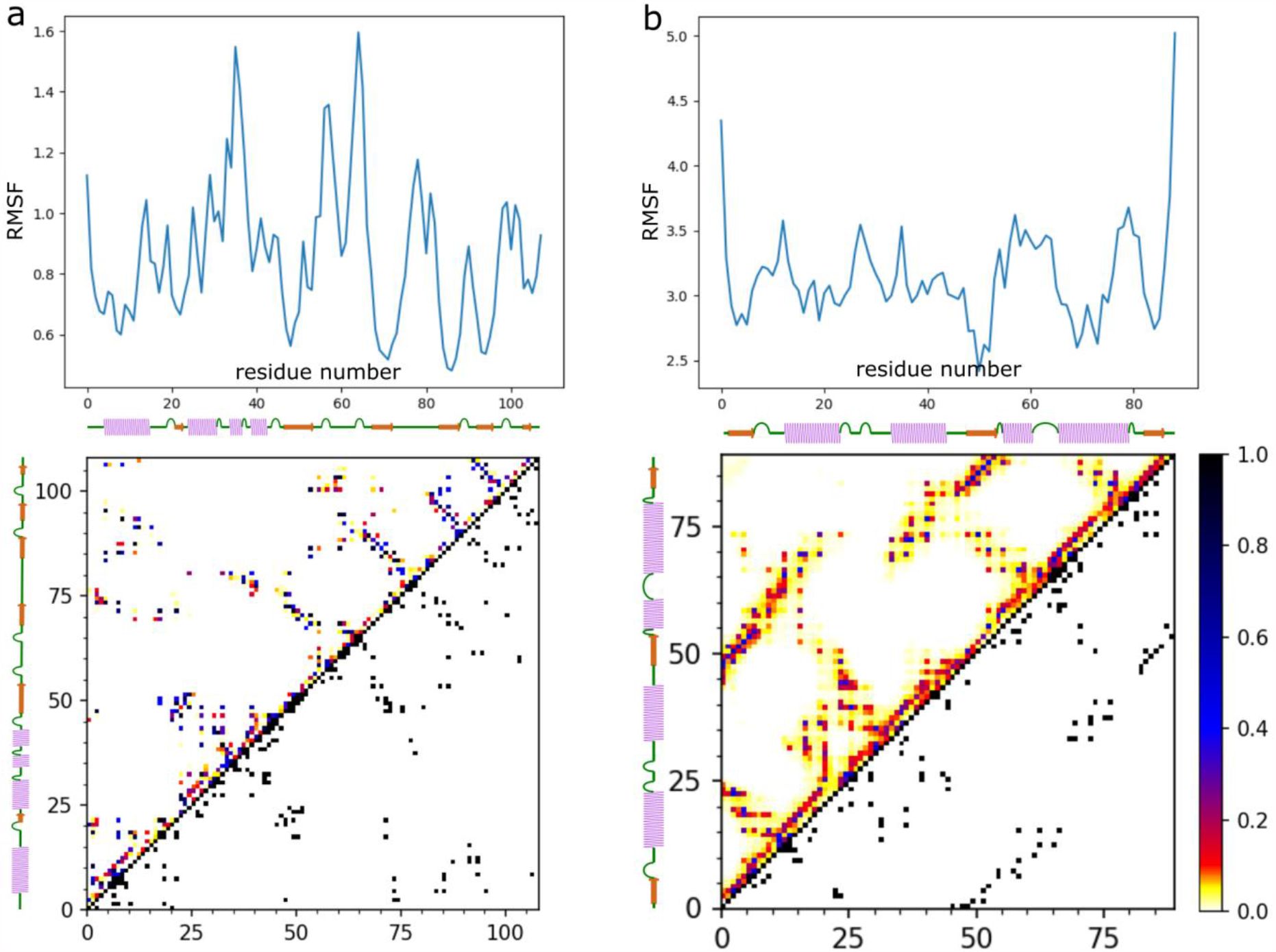
Characterization of barnase/barstar flexibility in the docking simulation. The figure shows RMSF plots (upper panels) and contact maps (lower panels) for (a) the barnase receptor and (b) the barstar ligand. The RMSF profile (see Methods) and contact maps showing the frequency of contacts are derived from the entire simulation (derived from 10’000 models).

**Figure 6.**
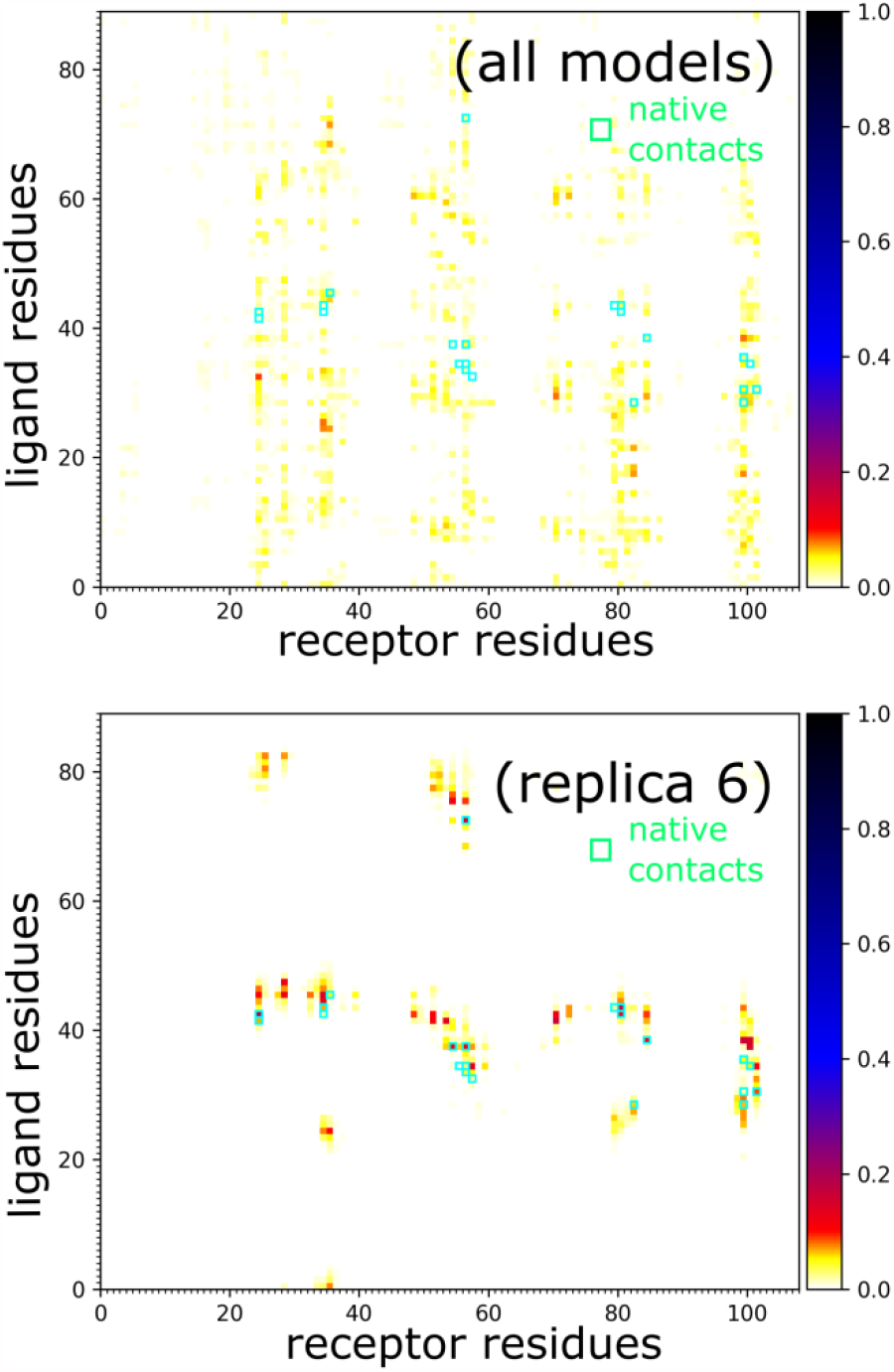
Characterization of barnase/barstar contacts. Panels show barnase/barstar models and contact maps for: (a) entire simulation (all, 10’000 models), (b) single selected replica (replica #6, 500 models) that reached a near-native arrangement. In the maps, green circles mark the native contacts.

An essential and unique feature of the presented docking simulations is the level of backbone flexibility during docking. In the example above, the ligand backbone fluctuations (LoRMSD) are in the range of 2-7 Å (Figure 4b), with the average LoRMSD value of 3.3 Å from the entire docking simulations. In other cases, the ligand fluctuations are at a similar level or higher (see LoRMSD values in Table 1).

Finally, using structural clustering of contact maps (see Methods), we attempted to select the set of 10 top-scored models for each case. Table 1 reports the most accurate models out of the 10 top-scored.

## Discussion

This work demonstrates a significant improvement in the sampling of large-scale conformational transitions during global protein-protein docking compared to other state-of-the-art approaches. We show that modeling the large conformational changes is possible at a relatively low computational cost. The presented simulations took between 10 and 80 hours (depending on the system size) using a single standard CPU. The proposed modeling protocol can be used as the docking engine in template-based and integrative docking protocols using experimental structural data and additional information from various sources [2,62]. We focused on the free docking of protein ligands with a highly flexible backbone in the present test simulations. Using unbound structures as the input, we produced acceptable accuracy models (iRMSD around 4 Å or lower) in low-flexibility and medium-flexibility cases. However, the selection procedure of the most accurate models needs further improvement. Namely, selecting the best-ranked models led to acceptable models in about half of the tested cases.

Presently, the most common approach to account for conformational changes in protein docking is using ENM [24–28,36–38]. The applicability of ENM to modeling protein flexibility is limited to specific systems and depends on how collective the protein motions are. Our method presents a conceptually different approach that seems to be more realistic (see review discussing coarse-grained CABS dynamics in the context of ENM approaches [24]). We demonstrated that it is possible to simulate effectively free docking of highly flexible protein ligands to quite elastic protein receptor structures. Such a significant degree of flexibility was achieved using a highly efficient simulation engine based on the coarse-grained representation of protein structures, Monte Carlo dynamics, and knowledge-based force field. CABS coarse-graining, enhanced by the discretized protein model and interaction patterns, significantly reduces the search space. Monte Carlo dynamics, enhanced by Replica Exchange annealing, leads to huge speed-up of the search procedures. Also a significant (although acceptable for many problems) flattening of energy surfaces by statistical potentials of CABS model simplifies simulations. As a result the flexible docking using CABS-dock is orders of magnitude faster than equivalent simulations based on classical modeling methods. Obviously the new method has also several limitations that have to be taken into consideration when designing new computational experiments. First, since the “ligand” protein is treated as a very elastic object (what is necessary to guarantee efficient search of the binding sites and poses) the cost of computations rapidly grows with the protein size. Thus, completely free global docking of protein ligands larger than 150 residues (see Table 1) may be impractical. Second, the coarse graining of the sampling space and simplification of interaction patterns (so important for the huge acceleration of the simulations) makes the docking energetics less sensitive. For these reasons the clustering procedures, refinement of the resulting structures and final model selection becomes very challenging and needs further developments. Also speeding-up the entire protocol can be useful. We estimate that the simulations could be easily speeded-up at least 10 times or more through algorithm parallelization. The speed-up would enable making the protocol available as the publicly accessible and automated web service.

## Conclusions

In summary, the described docking procedure accounts for large-scale protein structure fluctuations during unrestrained protein-protein docking search for the binding site. The exploration of such vast conformational space has not been demonstrated before to the best of our knowledge. The approach shows unprecedented sampling possibilities; however, the accuracy of the obtained complexes is still lower than observed for state-of-the-art docking tools. Definitely, the balancing of the structural restraints scheme needs further developments and tests. Therefore, this work is the first step towards a mature protein-protein docking tool. The next development steps would involve modifications of the distance restraints scheme, which allow for different degrees of flexibility for appropriate protein fragments (now the presented algorithm treats the entire protein-ligand as very flexible) and force-field improvements. The proposed approach is also very promising in the refinement applications when searching for the binding site is not needed, and only the protein-protein interface needs to be optimized.

## Acknowledgments

SK, MZ and AK acknowledge support by the National Science Centre (NCN), Poland [MAESTRO2014/14/A/ST6/00088]. MK acknowledges support by the National Science Centre (NCN), Poland [501/D112/66 GR-6271]. SK also acknowledges funding in part by the National Science Centre, Poland [UMO-2020/39/B/NZ2/01301].

